# Inference and design of antibody specificity: from experiments to models and back

**DOI:** 10.1101/2023.10.23.563570

**Authors:** Jorge Fernandez-de-Cossio-Diaz, Guido Uguzzoni, Kévin Ricard, Francesca Anselmi, Clément Nizak, Andrea Pagnani, Olivier Rivoire

**Author notes:** JFdCD and GU contributed equally to this work.

## Abstract

Exquisite binding specificity is essential for many protein functions but is difficult to engineer. Many biotechnological or biomedical applications require the discrimination of very similar ligands, which poses the challenge of designing protein sequences with highly specific binding profiles. Current methods for generating specific binders rely on *in vitro* selection experiments, but these have limitations in terms of library size and control over specificity profiles. We present a multi-stage approach that overcomes these limitations by combining high-throughput sequencing of phage display experiments with machine learning and biophysical modeling. Our models predict the binding profiles of antibodies against multiple ligands and generate antibody sequences with desired specificity profiles. The approach involves the identification of different binding modes, each associated with a particular ligand against which the antibodies are either selected or not. We demonstrate that the model successfully disentangles these modes, even when they are associated with chemically very similar ligands. Additionally, we demonstrate and validate experimentally the computational design of antibodies with customized specificity profiles, either with specific high affinity for a particular target ligand, or with cross-specificity for multiple target ligands. Overall, our results showcase the potential of leveraging a biophysical model learned from selections against multiple ligands to design proteins with tailored specificity, with applications to protein engineering extending beyond the design of antibodies.

## INTRODUCTION

Proteins often exhibit a delicate balance of multiple physical properties. A prominent example is binding specificity, where some ligand interactions are advantageous while others are detrimental. Examples include transcription factors, which recognize specific DNA motifs among a myriad of alternatives [1], enzymes with a strong preference for their substrate over many similar molecules [2, 3], and immune receptors capable of distinguishing a pathogenic molecule from many others, in particular self molecules [4]. Due to the close chemical similarity between favorable and unfavorable ligands, and/or the dissimilarities between favorable ligands, the engineering of such proteins poses formidable challenges. For instance, in the particular case of therapeutic antibodies, the desired specificity profile typically consists of strong binding affinity to the target antigen while retaining low binding affinity to human self antigens to avoid auto-immune reactions. Additionally, when the target antigen is a human protein, *e*.*g*. a tumor marker, antibody cross-specific binding to the human and the cyno and/or murine homologous antigens is often desired to ease drug development [5].

Presently, methods for obtaining specific binders essentially rely on *in vitro* selection experiments [6]. Phage or ribosome display [7, 8] with one immobilized targeted ligand in the presence of soluble non-targeted ligands allows screening for specific binding to the targeted ligand [9]. Yeast display combined with fluorescent-activated cell sorting [10] additionally offers the unique possibility to control precisely specificity selection criteria (including cross-specificity) upfront during the screening process by monitoring fluorescence associated with the targeted and non-targeted ligands in different channels [11], albeit with a maximum library size that is several orders of magnitude smaller. High-throughput selection can be combined with high-throughput sequencing read-out to identify binders beyond the top hits [12–14], but all experimental approaches are limited by the maximal library size, ranging from typically 10^8^ (yeast), 10^10^ (phage) to 10^15^ (ribosome). As large as these numbers may appear, they represent a negligible fraction of the combinatorially large space of possible sequences. Moreover, experimental screening for specificity requires the targeted and non-targeted ligands to be physically separable, which may be complicated if not impossible in some cases, for instance when considering distinct epitopes on the same molecule. Finally, in experiments, non-targeted ligands are inevitably present, since targeted ligands are typically attached to a cell, a tube/plate, or a magnetic bead.

Recently, works combining high-throughput sequencing and machine learning have demonstrated the possibility of making predictions beyond the scope of experimentally observed sequences [15, 16]. While past works predominantly focused on a single protein property (binding, stability, or catalysis) directly linked to the selection criterion [17], a few studies have shown the feasibility of inferring multiple physical properties, including quantities that are not directly measured [18]. Notable successful examples include predicting thermal stability from binding affinity measurements [19], and inferring specificity profiles of transcription factors from the selective enrichment of DNA sequences [20, 21]. Several recent works have started to apply this type of approach to predict and design antibody specificities [22–25] but, to our knowledge, none have addressed the critical but most challenging problem of designing antibody sequences that discriminate closely related ligands.

Here, we introduce a biophysics-informed approach that tackles this task. Trained on a set of experimentally selected antibodies, our model associates to each potential ligand a distinct binding mode, which enables the prediction and generation of specific variants beyond those observed in the experiments. To showcase this approach, we conducted a series of phage display experiments involving antibody selection against diverse combinations of closely related ligands. First, we demonstrate the model’s predictive power by using data from one ligand combination to predict outcomes for another. Second, we show its generative capabilities by using it to generate antibody variants not present in the initial library that are specific to a given combination of ligands. Our results highlight the potential of biophysical-informed models to identify and disentangle multiple binding modes associated with specific ligands. This approach has applications in designing antibodies with both specific and cross-specific properties and in mitigating experimental artifacts and biases in selection experiments.

## RESULTS

We designed phage display experiments for the selection of antibody libraries and performed two distinct experimental campaigns: in the first, we selected antibodies against various combinations of ligands. This provided us with multiple training and test sets, which we used to build and assess our computational model. In the second, we tested variants predicted by our model but not present in the training set to assess the model’s capacity to propose novel antibody sequences with customized specificity profiles.

### Experimental selection

Following our previously established protocols [13, 14], we carried out phage-display experiments with a minimal antibody library based on a single naïve human *V*_*H*_ domain in which four consecutive positions of the third complementary determining region (CDR3) are systematically varied to a large fraction of the 20^4^ = 1.6 10^5^ combinations of amino acids (“Germline library” [14]). This library is small enough to allow a high-coverage of its composition by high-throughput sequencing. Out of the 20^4^ potential variants, 48% are observed by sequencing, while we consider the remaining ones to be absent or unobserved. We previously showed that this library contains antibodies that bind specifically to a diversity of ligands, including proteins, DNA hairpins, and synthetic polymers [13, 14].

Here, we perform selections against complexes comprising two types of ligands, DNA hairpin loops and the surface of streptavidin-coated magnetic beads on which the DNA hairpins are immobilized. We performed independent selections against two such complexes, referred to as “Black” for one DNA hairpin on beads, and “Blue” for another DNA hairpin on beads, as well as selections against mixtures of Black and Blue complexes (“Mix”). Following standard protocols, we performed two rounds of selection with an amplification step in between, where each selection is preceded by an incubation of the phages with naked beads to (partly) deplete the antibody library of bead binders (see Fig. 4). These pre-selections provided us with data with a fourth selective pressure where naked beads are the only ligand (“Bead”). Importantly, we systematically rescued phages by infection of *E. coli* bacteria at each step of the protocol to closely monitor the antibody library composition. Input phages, phages bound to naked beads during the pre-selection step, and output phages bound to DNA target-coupled beads during the selection step were thus rescued in bacteria and extracted plasmids used as a template for high-throughput sequencing (see Fig. 4). The relationships between the 8 selection experiments are represented in Figure 1, together with the sequencing data that we collected.

**FIG. 1.**
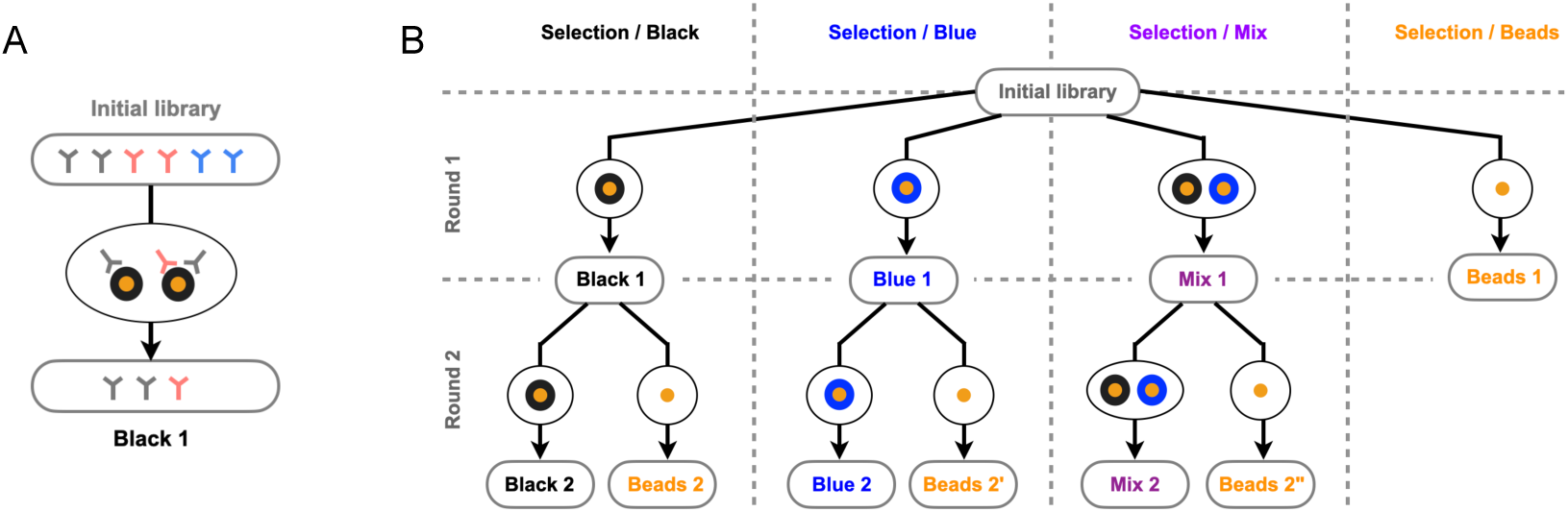
**A**. In a phage display experiment, an initial library containing ∼ 10^5^ variants, each in ∼ 10^6^ copies (here illustrated with 3 variants in 2 copies) is incubated in the presence of DNA hairpins (in black) coupled to magnetic beads (in orange). Antibodies are selected in proportion to their binding probability. The input and output populations are sampled and sequenced to provide data-sets of ∼ 10^5^ sequences each. **B**. We selected the same initial library against four different combinations of ligands: two different DNA hairpins coupled to magnetic beads, presented either alone or in combination, and naked magnetic beads. We refer to these four combinations as “Black”, “Blue”, “Mix” and “Beads” complexes. For the Black, Blue, and Mix complexes, we made two successive rounds of selection. The 10 boxes at the tip of the arrows indicate the 10 sequencing datasets thus produced to feed our model, in addition to the sequencing dataset from the initial library.

For each experimental selection round *t*, empirical enrichments were computed for each sequence *s* as *ϵ*_*st*_ = *R*_*st′*_ */R*_*st*_, where *R*_*st*_ and *R*_*st′*_, denote respectively the sequencing counts before and after selection. Enrichments against the Black and Blue complexes are observed to be very correlated, consistent with their close chemical similarities (Fig. 7). Enrichments against one complex and the beads alone are less correlated, indicating both that the beads are not dominant epitopes when coupled to DNA hairpins, and that they are chemically more dissimilar from these hairpins (Fig. 14).

### A model for multiple-specific selection

We built a computational model where the probability *p*_*st*_ for an antibody sequence *s* to be selected in a particular experiment *t* is expressed in terms of selected and unselected *modes*. Each mode *w* is mathematically described by two quantities: *μ*_*wt*_ that depends only on the experiment *t*, and *E*_*ws*_ that depends on the sequence, such that

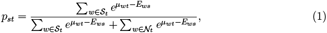

where *𝒮*_*t*_ and 𝒩_*t*_ represent, respectively, sets of selected and not-selected modes available in experiment *t*. (1) is rooted in the thermodynamics of binding at thermal equilibrium [26]: if a molecule can be in one of the selected (*𝒮*_*t*_) and unselected (*𝒩*_*t*_) thermodynamical states, its probability to be selected is given by (1), which corresponds to a Boltzmann law with *E*_*ws*_ = ∆*F*_*ws*_*/RT*, where ∆*F*_*ws*_ represents the free energy of *s* in state *w, R* the universal gas constant and *T* the temperature. A selected state can represent binding to a targeted ligand *w*, in which case *μ*_*wt*_ = ln[*w*], where [*w*] is the relative concentration of free ligand *w* in the experiment *t* (up to a scaling factor). The formula further includes an unselected unbound mode, to account for the possibility that the molecule remains in solution instead of binding any ligand.

Given that our experiments include three types of ligand – two DNA hairpins and magnetic beads – our model comprises four binding modes in total. A bead-bound mode is always selected, the DNA hairpin-bound modes are either selected or absent depending on the ligand present in the experiment, and the unbound mode is always unselected (Fig. 6). In addition to these physical modes associated with the thermodynamics of binding, our model can incorporate extra *pseudo* modes not related to binding, to account for biases that may occur during phage production and antibody expression stages (Materials and Methods for details). For each mode *w, E*_*ws*_ is parametrized by a shallow dense neural network (Materials and Methods). During training, the model parameters are optimized globally to capture the evolution of antibody populations across several experiments. Through this optimization process, the initial library abundances are also inferred (Materials and Methods). Once the model is trained, it can be used to simulate experiments with a custom set of selected/unselected modes, enabling the prediction of the expected probability of selection of variant reads, which can be compared to empirically observed enrichments of sequence counts in actual experiments.

Furthermore, we verified that introducing more complexity into the model along two different directions had a negligible impact. First, sequences recovered after one round of selection must be amplified before undergoing another round of selection, which occurs through phage infection and may be subject to biases. We collected sequencing data right before and after amplification and verified that no significant amplification bias was present (Fig. 9). Second, our model considers antibody sequences at the amino-acid level but selection can potentially occur at a nucleotidic level as well. We analyzed the data at this level and confirmed that no significant codon bias was observed in our experiments, consistent with an interpretation of the selection modes as arising primarily from ligand binding (Fig. 10).

### The model disentangles the effect of mixed ligands

To assess the model’s ability to disentangle the contribution to the selection of the different ligands, we conducted two types of validation.

#### Predicting selection against unseen mixtures of ligands

In the first validation, we trained the model using selection experiments against the Black and Blue complexes consisting of DNA hairpins attached to magnetic beads, and used the inferred model to predict the outcomes of experiments where these two complexes are mixed in equal proportions, which defines the Mix complex (Fig. 1). To assess the model’s performance, we compared the read counts of variants in the validation set with the abundances predicted by the model (Fig. 2A), and estimated the correlation between predicted probabilities of selection *p*_*st*_ and experimentally determined enrichments _*st*_ against Mix (Table I). The results validate the model’s capacity to integrate different selection experiments to predict the results of selection experiments with unseen combinations and proportions of ligands. As a control, selectivities predicted using only one mode result in significantly lower correlations, confirming that both Black and Blue modes are necessary to model selection in the Mix experiment (Table I).

**FIG. 2.**
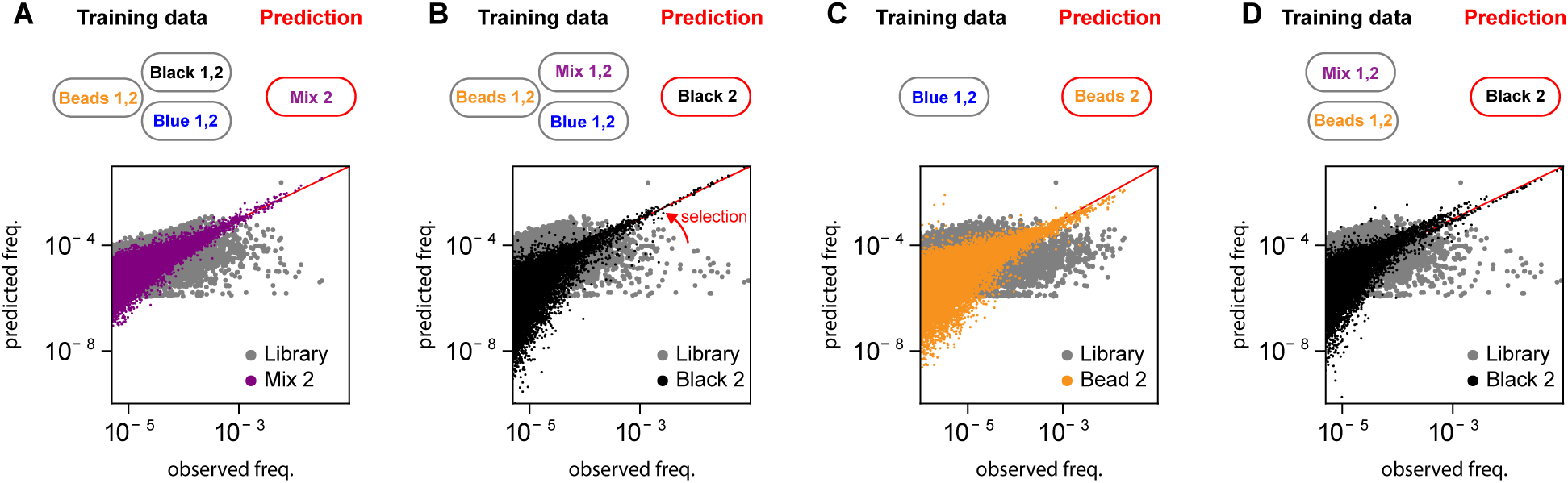
The model predicts accurately the evolution of sequence variants abundances in response to multiple selective pressures. We considered different tasks of increasing difficulty, depending on the training set used: **A**. Model trained on the experiments with Black, Blue complexes, and empty Beads, and prediction evaluated with a mixture of the Black and Blue complexes; **B**. Model trained on experiments with a mix of Black and Blue complexes, Blue complexes only, and naked Beads, with predictions evaluated on the experiment with Black complexes only; **C**. Model trained on experiments with Blue complexes only, and predictions evaluated on experiments with naked Beads; **D**. Model trained on experiments with mix of of Black and Blue complexes and naked Beads, and predictions evaluated on experiments with Black complexes only. The panels show scatter plots of the the observed (*x*-axis) vs. predicted sequence frequencies (*y*-axis), with the initial library abundances shown in gray for comparison. The correlation between empirical selectivities and the model-predicted selectivities for each task are given in Table I.

#### Predicting selection against hidden ligands

In the second validation phase, we trained the model to predict selections against unseen subsets of ligand combinations. We considered three scenarios of increasing complexity: (i) using the data from the Mix, Beads, and Blue selections to disentangle the Black mode and predict the experiment with the Black complex (Fig. 2B), (ii) disentangle the effect of Beads using Blue data exclusively and predict the Beads selection (Fig. 2C), and (iii) disentangle the Black ligand effect from Mix and Beads selections and predict the Black selection experiment (Fig. 2D). The second task is more challenging than the first because the beads in the Blue complex are subdominant epitopes (Fig. 14), and the third task is more challenging than the other two because the two hairpins are very similar (Fig. 7) and not seen independently.

As previously, we compared in each case predicted selectivities to empirical enrichments from experiments and obtained very good correlations (Fig. 2 and Table I). Altogether, these results validate the model’s capacity to disentangle the contributions of different ligands, and effectively “subtract” the contribution of some ligands to predict the contribution of others.

### Generation and validation of antibodies with custom specificity profiles

In addition to predicting the outcome of experiments involving new combinations of ligands, our model can be employed to design novel antibody sequences with predefined binding profiles. These profiles can be either cross-specific, allowing interaction with several distinct ligands, or specific, enabling interaction with a single ligand while excluding others. The generation of new sequences relies on optimizing over *s* the energy functions *E*_*sw*_ associated with each mode *w* in (1). To obtain cross-specific sequences, we jointly minimize the functions *E*_*sw*_ associated with the desired ligand. On the contrary, to obtain specific sequences, we minimize *E*_*sw*_ associated with the desired ligand *w* and maximize the ones associated with undesired ligands.

Panel A of Figure 3 illustrates the distribution of sequences in the energy plane defined by the modes associated with the two DNA hairpins, as inferred when using all available data. Among all possible sequences, we select those not present in the initial library (thus not included in the training set) and predicted to possess specific binding profiles: sequences in blue are predicted to bind strongly to the Blue DNA hairpin while exhibiting weak binding to the Black DNA hairpin, and reciprocally for those in black, while those in purple are predicted to bind both hairpins.

**FIG. 3.**
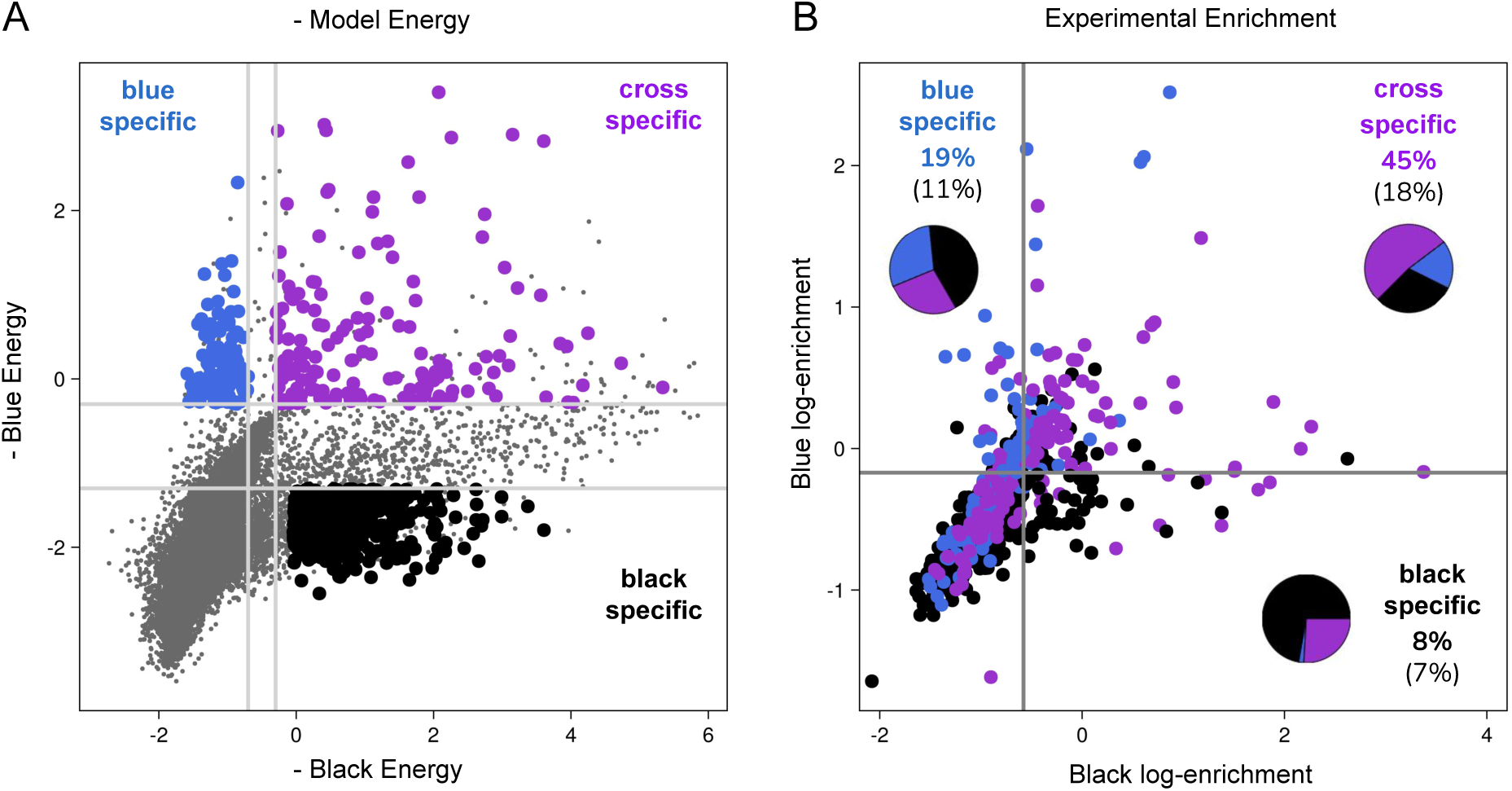
Design and validation of antibodies with prescribed specificity. **A**. Model-based energy plot where each sequence *s* is represented as a circle with coordinates 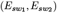, with *w*_1_ representing the binding mode associated with the Black hairpin and *w*_2_ with the Blue hairpin. Sequences predicted to be specific to the Blue hairpin, specific to the Black hairpin, or cross-specific to the two hairpins are respectively highlighted in blue, black, and purple. We selected for experimental validation all the colored sequences that are not present in the training set. **B**. Experiment-based enrichment plot of the selected sequences where each sequence *s* is represented as a circle with coordinates 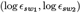, with 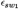 representing the enrichment against the Black complex and *w*_2_ against the Blue complex. Sequences with high enrichment in one experiment and low enrichment in the other are ligand-specific, those with high enrichment in both are cross-specific, and low-enrichment sequences are non-binders (false positives). We assess our computational approach’s effectiveness by calculating the percentage of designed sequences falling within the correct region. Thresholds are set based on the average enrichment of all sequences in the experiment including the control sequences (see Fig. 11 for more details). Cross-specific designed antibodies achieve a 45% true positive rate, while Black and Blue-specific binders yield lower percentages (19% and 8%, respectively), reflecting the capacity of our approach to design antibodies with desired properties despite the challenges arising from the close similarity of the two ligands.

We validated experimentally these predictions by phage display selection of a library composed of these ∼ 2000 computationally designed sequences and ∼ 10 control sequences for binding to either Black or Blue complexes. Panel B of Figure 3 provides a summary of the results. The enrichments of variants in the two experiments are displayed, with variants above two predefined thresholds (see Fig. 11 for details) considered as binders. The four regions represent specific binders for Black and Blue DNA hairpins, cross-specific binders, and non-binding variants. Percentages of the designed antibodies that fall within the respective regions (true positives) are reported, along with the fraction of the total number of points for comparison. Additionally, the composition of variants within the region segmented by designed specificity is presented. Taken together, these results demonstrate the capacity of the model to propose multiple sequences with desired specificities. Not all designed antibodies have the desired properties, but it must be stressed that the results of Figure 3 address the case of two very similar ligands with the further constraint that the initial library already covers half of the potential diversity, which leaves a relatively small novel design space. In contrast, designing binders to a single ligand regardless of their affinity to the others is comparatively easier (Fig. 12).

## DISCUSSION

In this study, we propose a multi-stage method that combines high-throughput sequencing of coordinated phage-display experiments, with a machine-learning approach that trains a biophysical model. This model is designed to capture statistical patterns associated with different aspects of the selective pressures to which antibodies are experimentally subjected. By disentangling the different factors influencing selection, we can design sequences with novel combinations of physical properties, making the most of the wealth of information contained in high-throughput sequencing data from selection experiments.

Over the past three years, several machine learning approaches have been developed with the aim of analyzing antibody selection experiments to propose new antibody variants with improved binding affinities for a prescribed target, given particular constraints. These constraints include parameters such as viscosity, clearance, solubility and immunogenicity [23], which are important for drug development, or non-specific binding [24], to eliminate antibodies that tend to bind indiscriminately to a large number of antigens. Some of these works are based on experimental data similar to ours, combining selections against multiple targets with a similar aim of extracting target-specific features [22].

Our work differs from these studies in the difficulty of the task we are tackling. We focus indeed on inferring and designing high levels of binding specificity, which involves discriminating between molecular targets with significant structural and chemical similarity. To provide a clear proof of concept, we considered two targets that are not of direct biomedical interest but whose similarity is well characterized. Our two 24-nucleotide hairpins thus differ only by 7 nucleotides in their loop. This difference is comparable to the difference between DNA sequences recognized by transcription factors or restriction enzymes, some of the most specific proteins found naturally. Generating data and developing a model from which to design sequences that discriminate between these two targets is a very rigorous test.

A practical application of our approach is the design of new protein candidates with prescribed specificity profiles. The minimal breadth of our initial library reduces the possibility of testing entirely new variants, but our approach is also applicable to libraries of greater breadth. As these libraries are necessarily much more undersampled, the potential for discovering better variants is greater, although undersampling can also lead to less accurate models. Finally, although not all the variants proposed by our model proved experimentally to have the desired properties, a significant fraction did, which is enough to provide several alternative sequences at the typical scale of ∼100 variants that can be tested experimentally.

There are several avenues for extending the scope of our work. One is to increase the diversity of the initial library, which also allows the incorporation of additional physical modes associated, for example, with thermal stability. Another is to generate and integrate data from experiments in which ligand libraries are selected to bind to one, or several, binders. Beyond practical applications, these extensions have the potential to provide a general approach to deducing the multiple physical properties encoded in protein sequences.

## MATERIALS AND METHODS

### Phage display selection

Phage display selections were performed essentially as in our previous study [14]. Our ‘Germline’ *V*_*H*_ library [14] and the library of designed sequences are both cloned in the pIT2 phagemid vector. M13KO7 (Invitrogen) was used as a helper phage, and TG1 *E. coli* as a host. M280 streptavidin-coated magnetic beads (Dynal) were used for DNA target immobilization. DNA targets are single-stranded DNA oligonucleotides biotinylated at their 5’ end (IDT). For the selection against Mix, beads coupled to the Black DNA hairpin target were mixed 50-50 with beads coupled to the Blue DNA hairpin target.

Phage display experiments included a pre-selection step with naked beads followed by a selection step with DNA target-coupled beads. Specific to the present study, we rescued phages at three steps of the selection process (see Fig. 4), namely (i) input phages, (ii) output phages bound to naked beads during pre-selection, (iii) output phages bound to DNA target-coupled beads during selection. The exact same washing and elution procedures were applied to naked beads and DNA target-coupled beads prior to phage rescue in TG1 cells. Consistent with efficient selection for DNA target-binding, we typically observed a 10 to 100-fold higher phage titer in elutions from beads coupled to DNA targets (10^6^ to 10^7^ phages) than from naked beads (10^5^ phages).

### Sequencing read-out

For each phage sample to be sequenced, an amplicon encompassing the 4 randomized CDR3 sites flanked with Illumina adapters bearing sample-specific indices was produced by PCR on DNA purified from TG1 cells following phage rescue. The number of PCR cycles was kept as low as possible to avoid distortion due to amplification biases, which we checked specifically.

The ‘Germline’ library selection was sequenced on the Illumina™ NextSeq 500 platform, producing 76 bp reads. The in-silico designed library selection was sequenced on the Illumina™ NextSeq 1000 producing 60+60 bp (paired ends) reads.

### Model training

The model is trained by maximizing the likelihood of the observed sequencing read counts of each sequence *s* in an experiment *t*, that we denote by *R*_*st*_, and which are modeled as a multinomial distribution:

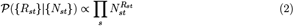

where *N*_*st*_ is the estimated abundance of this variant in the experiment. The abundances evolve from one experiment *t* to the next *t*^*′*^, according to *N*_*st′*_ ∝ *p*_*st*_*N*_*st*_, where *p*_*st*_ are the selectivities in (1). Iterating this relation, we can express *N*_*st*_ as a function of the abundances in the initial library, *N*_*s*0_. Since the *N*_*s*0_ are not directly observed, they are also inferred by maximum likelihood. The Mathematical Supplement contains more details about the model and its implementation.

An *L*_2_ squared norm regularization is added to penalize large fields (in the independent-site model), or the neural network weights. Training is substantially accelerated by splitting the sequences into mini-batches. In practice, we form random batches of 128 sequences, which are re-shuffled at each epoch. Further details are given in Supplementary Materials.

### Processing of the data

Sequences containing stop codons are discarded. They are either sequencing errors or can be enriched during amplification since the expression of the antibody is costly for the bacteria. To further reduce sequencing errors, we sequences where the flanking constant regions of the CDR3 do not coincide with the designed framework sequence are also fitered out.

### Data and Software Availability

All sequencing data generated from our selection experiments will be shared on the Sequence Read Archive (SRA), the primary NIH-funded archive for high throughput datasets. This data received the accession code BioProject ID PRJNA1028404 and can be accessed at http://www.ncbi.nlm.nih.gov/bioproject/1028404. The code to reproduce the results in this paper is available at https://github.com/2023ab4/ab4.

## ACKNOWLEDGMENTS

AP, GU, and JFdCD acknowledge funding from European Unions Horizon 2020 research and innovation programme MSCA-RISE-2016 under grant agreement No.734439 INFERNET. CN acknowledges support from ANR-17-CE30-0021. AP acknowledges financial support from Future Artificial Intelligence Research (FAIR) and Centro Nazionale di Ricerca in High-Performance Computing, Big Data, and Quantum Computing (ICSC) founded by the European Union-NextGenerationEU. OR acknowledges support from FRM AJE20160635870 and ANR-21-CE45-0033. We thank Carlo Cosimo Campa for interesting discussions on sequencing strategies.

## Appendix A: Detailed experimental methods

### Phage display selection

Phage display selections were performed essentially as in our previous study [14] with our ‘Germline’ *V*_*H*_ library [14] or the library of designed sequences, both cloned in the pIT2 phagemid vector. M13KO7 (Invitrogen) was used as a helper phage, and TG1 *E. coli* (Agilent) as a host. Beads coupled to DNA were prepared by adding 10μL of 400μM biotinylated ssDNA target (IDT) incubated for 15mn with 50μL M280 streptavidin-coated magnetic beads (Dynal) that had been washed prior according to the manufacturer’s instructions, followed by two additional washing steps. The Black biotinylated ssDNA target sequence is biotin-AAAAGACCCCATAGCGGTCTGCGT. The Blue biotinylated ssDNA target sequence is biotin-AAAAGACTTGGTAATAGTCTGCGT. Both ssDNA targets form a hairpin sharing a common stem and bearing different loops. For the selection against Mix, beads coupled to the Black DNA hairpin target were mixed 50-50 with beads coupled to the Blue DNA hairpin target.

The general scheme of our phage display experiments is described in Fig. 4. Experiments included a pre-selection step with naked beads followed by a selection step with DNA target-coupled beads. Specific to the present study, we rescued phages at three steps of the selection process, namely (i) input phages, (ii) output phages bound to naked beads during preselection, (iii) output phages bound to DNA target-coupled beads during selection. The exact same washing and elution procedures were applied to naked beads and DNA target-coupled beads prior to phage rescue in TG1 cells.

Input phages were produced for 7h at 30°C following infection with a 20-fold excess of helper phages, and the culture supernatant used as is (no phage precipitation to avoid phage clusters). Our libraries were screened by pre-selection of 10^11^ input phages in 1mL against 50μL naked magnetic beads for 90mn, followed by selection of phages that were not bound to naked beads against biotinylated target DNA hairpins immobilized on 50μL magnetic beads for 90mn. Ten washing steps were performed with 8mL 0.1% Tween20 (Sigma) prior to elution with 1mL 100mM triethylamine (Sigma) for 20mn and neutralization with 0.5 mL Tris 1M pH=7.4 (Sigma), both on naked beads to recover bead binders and on DNA target-coupled beads to recover DNA target binders via phage rescue by infection of an excess of TG1 cells. Input phages were also rescued in TG1 cells by adding 500μL TG1 to 500μL of a 100-fold dilution of the input phage stock.

Consistent with efficient selection for DNA target-binding, we typically observed a 10 to 100-fold higher phage titer in elutions from beads coupled to DNA targets (10^6^ to 10^7^ phages) than from naked beads (10^5^ phages).

#### Cloning of designed sequences

The library of ≈ 2000 designed sequences was constructed by PCR-amplification with the Q5 High-fidelity 2x Master mix (New England Biolabs) of a 120bp oligo pool (Twist Bioscience), the sequence of which encompasses the 4 randomized CDR3 sites, followed by homology-based assembly (HiFi NEBuilder, New England Biolabs) cloning into the same pIT2 vector as our ‘Germline’ *V*_*H*_ library.

#### Sequencing read-out

TG1 cells used for phage rescue (input, output from naked beads, output from DNA target-coupled beads), grown on solid plates overnight, were scraped and DNA extracted using a miniprep kit (Macherey-Nagel). A 200bp amplicon encompassing the 4 randomized CDR3 sites flanked with Illumina adapters bearing sample-specific indices was produced in 2 PCR steps. A first PCR uses purified pIT2 plasmid from TG1 cells as a template and staggering oligonucleotides (to favor clustering, as the upstream and downstream flanking sequences of CDR3 sites are constant) to add half of the Illumina adapters without indices. A specific pair of staggering oligonucleotides is used for every subsample to be sequenced (input, output with target or without target). A second PCR uses the product of the first PCR step as template and adds the indices and the remaining part of Illumina adapters. Both PCR steps were carried out with the Q5 High-fidelity 2x Master mix (New England Biolabs) for 15 cycles to avoid distortion due to amplification biases, which we checked specifically.

## Appendix B Analysis of empirical enrichments

For each experimental selection *t*, empirical enrichments were computed for each sequence *s* as

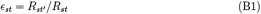

where *R*_*st*_ and *R*_*st′*_, denote respectively the sequencing counts before and after selection. Since the empirical estimate (B1) is susceptible to sampling noise, and can be undefined if *R*_*st*_ vanishes, in practice ϵ_*st*_ is computed only for sequences *s* for which both *R*_*st′*_ and *R*_*st*_ are larger than a minimum threshold count. As the diversity of the population decreases between the first and second rounds of selection, it is expected that these enrichments are best estimated at the second round of selection, although for a smaller set of sequences that are more represented.

Fig. 7 compares the enrichments obtained in different experiments, where selection corresponds to binding different targets. Fig. 8 plots the Pearson correlations between these enrichments, as a function of the minimum count threshold imposed in the numerator and denominator of (B1). Empirical enrichment against the Blue, Black, and Mix complexes show significant correlations (Fig. 8), consistent with the structural and chemical similarity of the two DNA hairpins. This feature reflects our choice to study closely related ligands that are challenging to differentiate. In contrast, empirical enrichments from selection against the Beads are appreciably more distinct from the other ones (bottom rows of Figures 7 and 8).

For variants with more than 50 counts before and after selection, we compute their empirical enrichment and compared this to the model predicted selectivity. Results are shown in Table I. We did this for each of the computational predictions in Fig. 2, and report here the resulting correlations in each case. As a control the last column of Table I reports the correlation if the mode Blue is used to predict the enrichments, instead of the correct one.

## Appendix C Mathematical supplement

### Biophysical model of selection

Let *N*_*st*_ be the total number of phages carrying sequence *s* in a library at round *t*, before selection. In one experimental round, phages are selected by some phenotypic criteria (*e*.*g*. binding to a target). In our model, we consider each sequence to be in one of several possible thermodynamic states *w* (target bound, unfolded, in solution, etc.). Each thermodynamic state *w* is populated by a number of particles *n*_*swt*_, and we have Σ_*w*_ *n*_*swt*_ = *N*_*st*_. Also, each thermodynamic state is described by a sequence-dependent energy function *E*_*sw*_, related to the propensity of a sequence to be found in this state (in the following we describe some possible parametrization of this function). We denote by *μ*_*wt*_ the chemical potentials for each state at a particular round, we can then model the abundances *n*_*swt*_ as follows,

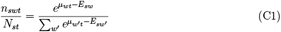

In the case of a thermodynamic state corresponding to binding a target, the chemical potentials *μ*_*wt*_ are proportional to the logarithmic concentration of the available target to bind. Next, we define the selectivity of a sequence *s* in the experimental round *t*, as:

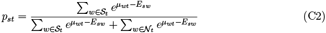

Here, we consider a subset of states 𝒮_*t*_ which are *selected* in the current experiment (*e*.*g*. bound to target), and a set of states *𝒩*_*t*_ which are depleted (*e*.*g*., washed in solution). Together, *𝒮*_*t*_ ∪ *𝒩*_*t*_ defines the set of feasible thermodynamic states in the experimental conditions of round *t*. Particles that adopt states *w* ∈ *𝒮*_*t*_, are selected and result in an enrichment of the corresponding sequence. The remaining (1 − *p*_*st*_)*N*_*st*_ particles of sequence *s* are washed away.

#### Amplification

After the selection step, there is an amplification step when the overall population size is restored. Assuming that the abundances are normalized, Σ_*s*_ *N*_*st*_ = 1 and denoting by *N*_*s,t*+1_ the phage abundances prepared for the next round, we have:

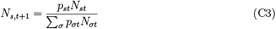

Here, (Σ _*σ*_ *p*_*σt*_*N*_*σt*_)^−*1*^ is the amplification factor necessary to restore the original population size after selection in round *t*.

#### Multiple rounds

When dealing with multiple selection experiments, the output of a selection round can be used as the input of another selection round. We consider the selection experiments to be arranged into a rooted tree, such as the one depicted in (Fig. 1). The nodes represent phage populations at a specific time with the root representing the initial sequence library. The edges represent a selection and amplification process that modifies the population in the parent node to the descendant node. In each branch, the node’s generations are denoted by *t*, with *t* = 0 being the root. In particular, *N*_*s*0_ represents the initial library abundances. The parent of a non-root node, *t >* 0, is denoted by *a*(*t*). Starting from a node *t*, we can traverse back to its ancestors until we reach the root of the tree. We define by:

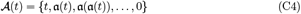

the set of ancestors encountered during this traversal (note that 𝒜 (*t*) includes *t* itself, for convenience). In particular, for the root node *𝒜* (0) = *{*0*}*. For *t >* 0, we have that *p*_*st*_ denotes the selectivity of sequence *s*, in the round of selection that brings the population from *a*(*t*) to node *t*. It follows that we can write:

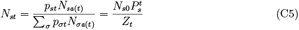

for *t >* 0, by induction, where 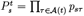 and 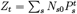. For the root nodes it is convenient to set *p*_*s*0_ = 1, *P*_*s*0_ = 1, *Z*_0_ = 1, and *a*(0) = 0. Then these formulas remain valid at the root. Given the selectivities *p*_*st*_ and the initial abundances *N*_*s*0_, we can use these expressions to determine all future populations of the tree.

#### Training the model

The data are the counts of sequence reads taken at each sequenced round (that could be a subset of all nodes in the experiment tree), *{R*_*st*_*}*. As the result of a sampling and sequencing procedure, the counts are related to the actual abundances by a multinomial distribution:

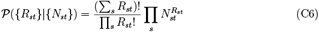

The abundances of all populations *{N*_*st*_*}* are determined by the initial abundances *{N*_*s*0_*}* and the selectivities, *{p*_*st*_*}*. Therefore, we can write *𝒫* (*{R*_*st*_*}*|*{N*_*st*_*}*) = *𝒫* (*{R*_*st*_*}*|*{p*_*st*_*}, {N*_*s*0_*}*). Since the initial abundances *{N*_*s*0_*}* are usually unknown, we can train our model by maximizing *𝒫* (*{R*_*st*_*}*|*{p*_*st*_*}, {N*_*s*0_*}*) with respect to the parameters determining the selectivities *{p*_*st*_*}* (to be specified below) and the initial abundances *{N*_*s*0_*}*, subject to *N*_*s*0_ ≥ 0 and Σ _*s*_ *N*_*s*0_ = 1. To carry out the maximization over *{N*_*s*0_*}* we can introduce a Lagrangian,

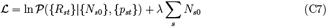

where we used the Lagrange multiplier *λ* for the constraint Σ _*s*_ *N*_*s*0_ = 1. Differentiating with respect to *N*_*s*0_ and setting the derivative to zero, gives the equation:

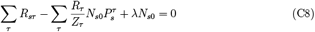

where *R*_*τ*_ = Σ _*σ*_ *R*_*στ*_, and *R*_*τ*_ */Z*_*τ*_ is the sampling coverage at round *τ* . Note that:

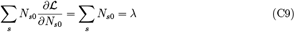

Therefore the stationarity conditions 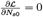imply *λ* = 0. This is also intuitively expected, because *ℒ* depends only on the relative ratios among the *N*_*s*0_, and would be insensitive to a global increase of the total Σ _*s*_ *N*_*s*0_ while preserving those ratios. Therefore *ℒ* has no gradient orthogonal to the constraint Σ _*s*_ *N*_*s*0_ = 1, making the Lagrange multiplier unnecessary. Now solving for *N*_*s*0_, we obtain:

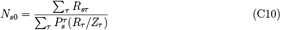

which gives *N*_*s*0_ as functions of *Z*_*t*_ and the selectivities *p*_*st*_. In particular note that *N*_*s*0_ = 0 for unobserved sequences (those for which *R*_*st*_ = 0 for all *t* in the tree). There are two contributions to the initial abundances: the different sampling coverage at different rounds and the effect of selection. In absence of selection, the formula above becomes more intuitive. Then *P*_*st*_*/Z*_*t*_ is a constant independent of *s, t*, which must be one by normalization. Then, *N*_*s*0_ = Σ _*τ*_ *R*_*sτ*_ */ Σ* _*στ*_ *R*_*στ*_ is also independent of *t*, and we just aggregate all the read samples to make an inference of the underlying abundances. The factors *P*_*sτ*_ *R*_*τ*_ */Z*_*τ*_ in the denominator of (C10), then serve to account for the effect of selection.

By using Equation (C10) we can consider *Z*_*t*_ as free parameters, in place of the initial abundances *N*_*s*0_. Obtaining the following learning gradient:

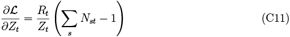

Notice that *ℒ* as a function of 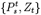 depends only on the ratios *Z*_*t*_*/Z*_*t ′*_ and is invariant to a multiplication of all the *Z*_*t*_ by a common factor. Therefore we can set *Z*_0_ = 1 to break this degeneracy, consistent with the previous definitions. The stationary conditions 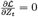reproduce the normalization constraints Σ _*s*_ *N*_*st*_ = 1. It follows that if we treat the *Z*_*t*_ as free parameters, at a stationary point of *ℒ* these constraints will be satisfied automatically.

#### Low-selectivity regime (or *rare binding* approximation)

A further simplification can be obtained by assuming a low-selectivity regime, where *p*_*st*_ « 1 for all sequences in all rounds. We can then approximate:

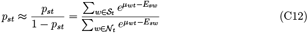

We call this the *rare binding approximation* (RBA) [16].

#### Sequence to energy mapping

The selectivities *p*_*st*_ are given by Equation (C2). In turn, a mapping giving the energies *E*_*sw*_ for each sequence must be parameterized. We here considered two models. In the simplest case, all the sites of the sequence are independent, which leads to an additive model,

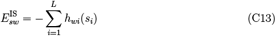

that we call the independent-site model (IS), and where the local fields *h*_*wi*_ are learned during model training. This assumption fails to consider epistatic effects between pairs of sites within a mode. To account for these, a possibility is to introduce a Potts-like model with two-body interactions, as typically done in DCA-like approaches [27, 28]. However this results in a large number of coupling parameters (∼ 20^2^*L*^2^). More generally, we can consider any functions *E*_*sw*_ = *E*_*w*_(*s*) assigning energies to sequences in different thermodynamic states. We here considered a feed-forward neural network, taking as input a one-hot encoded sequence, and producing as output the energy, 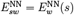. The parameters of the neural network (NN) are then learned during model training. Epistatic interactions arise as non-trivial correlations induced by the non-linearities of the network. We report the details of our architectural choices below.

#### Gauge invariance in the rare-binding approximation (RBA)

In the low-selectivity regime, a new gauge invariance appears. We make the following change of variables:

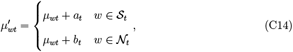

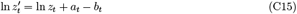

Then, under the RBA regime,

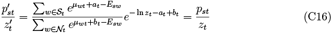

Thus, the amplification factors become indistinguishable from the chemical potentials. In other words: we cannot infer the amplification factors.

From the previous section recall that we imposed the gauge Σ _*w*_ *μ*_*wt*_ = 0. This means that *a*_*t*_, *b*_*t*_ are not completely free, but rather they must satisfy | *𝒮* _*t*_|*a*_*t*_ + | *𝒩* _*t*_|*b*_*t*_ = 0. The remaining degree of freedom can be exploited to enforce 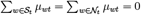, which is stronger than the condition Σ _*w*_ *μ*_*wt*_ = 0 from the previous section. More precisely, given chemical potentials *μ*_*wt*_, we can choose:

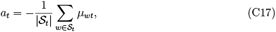

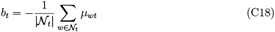

which satisfy |*𝒮*_*t*_|*a*_*t*_ + |*𝒩*_*t*_|*b*_*t*_ = 0, and for which:

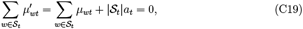

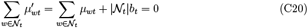

while the amplification factors transform as:

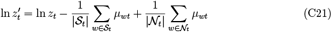

When the chemical potentials satisfy 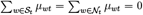, we will say that we are in the *rare binding gauge*.

This change of variables shows how one can absorb the amplification factors into the chemical potentials. Alternatively, we can exploit this new gauge invariance to impose that ζ_*t*_ = 0, by choosing 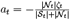 and 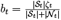 .

#### Architecture of feed-forward neural network

For each state considered, the architecture consists of 3 layers, with 20 and 5 hidden units, with a SeLU nonlinearity [29] in each.

#### Regularization

For the independent-site model, the regularization penalty has the form:

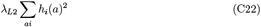

while for the NN model the sum extends over all weight parameters in the network. The coefficient *λ*_*L*2_ was panned over a range between 10^−4^ and 10. The optimal choice in terms log-likelihood of a withheld dataset was chosen, resulting in *λ*_*L*2_ = 0.01.

## Appendix D: Supplementary Figures

**FIG. 4.**
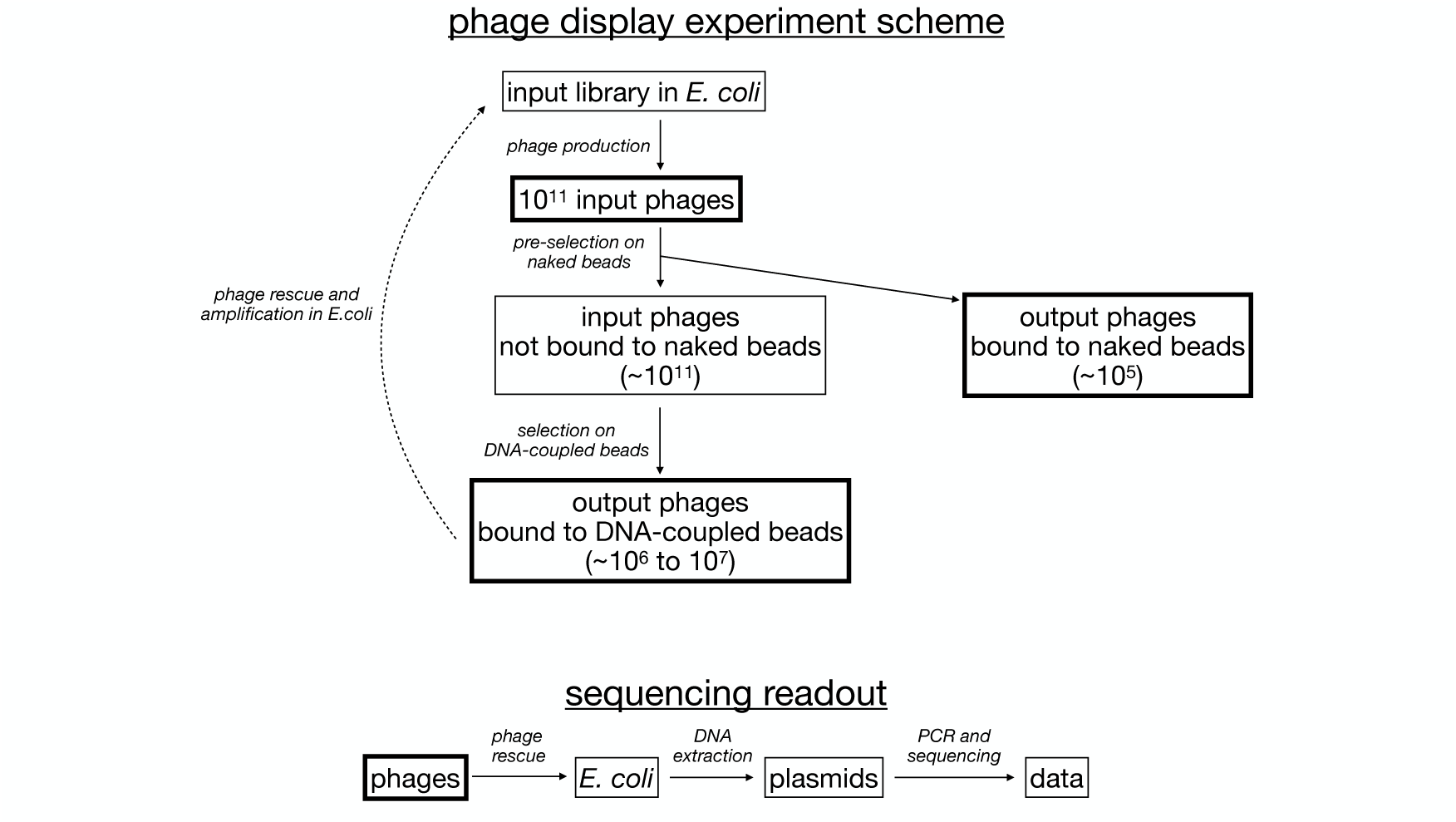
**Phage display experiment scheme**. Input phages are produced from *E. coli* bacteria and first pre-selected against naked beads. Phages that did not bind naked beads are then selected for binding to DNA target-coupled beads. Phages bound to DNA-coupled beads are rescued by infection of *E. coli* bacteria for the next cycle. **Sequencing readout**. Input phages, output phages bound to naked beads and output phages bound to DNA-coupled beads are rescued by infection in *E. coli* bacteria. Plasmid DNA is extracted from bacteria to serve as a template for high-throughput sequencing of a PCR amplicon encompassing the 4 randomized antibody CDR3 sites.

**FIG. 5.**
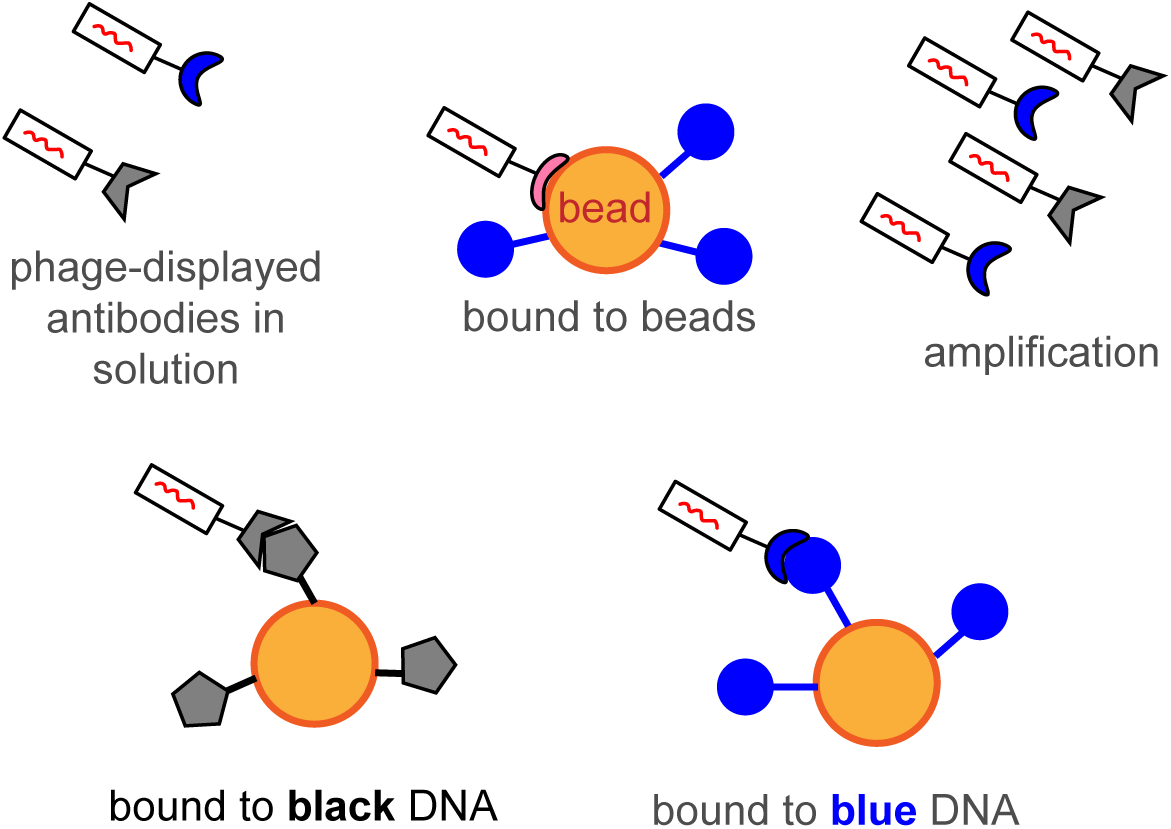
The figure illustrates the modes integrated into the model, each associated with distinct states: The first mode corresponds to an unselected, unbound state, and it is a common element in every selection round. The other modes are linked to the binding of either the Black and Blue ligand or the Beads, where they are immobilized. An additional mode exists in the model, which isn’t directly related to the physical binding to specific ligands, instead it accounts for the broader process of amplification and the potential biases introduced within the phage population. The selection or exclusion of these modes depends on the specific selection round, as visually depicted in Fig. 6.

**TABLE 1.**
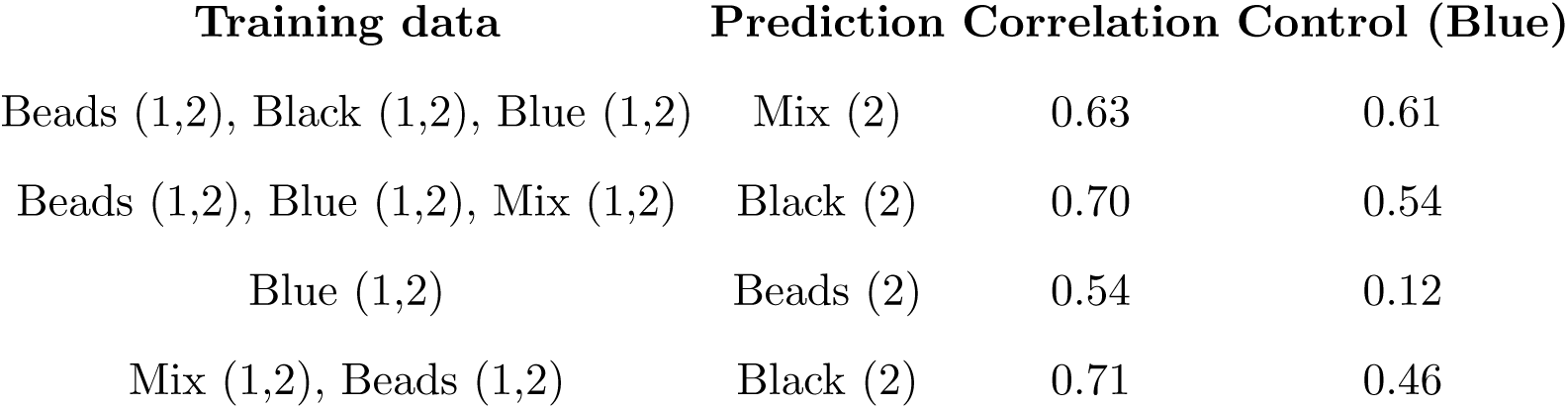
Correlations between predicted and empirical enrichments. A model trained on the data from the first column is used to predict enrichments of the experiment in the second column. The correlations between empirical log-enrichments and model log-selectivities are given in the third column. To demonstrate that the inferred modes have a meaningful correspondence to the physical modes, the last column shows (as a control) how the correlation decreases if only the Blue mode is used to predict the selectivity.

**FIG. 6.**
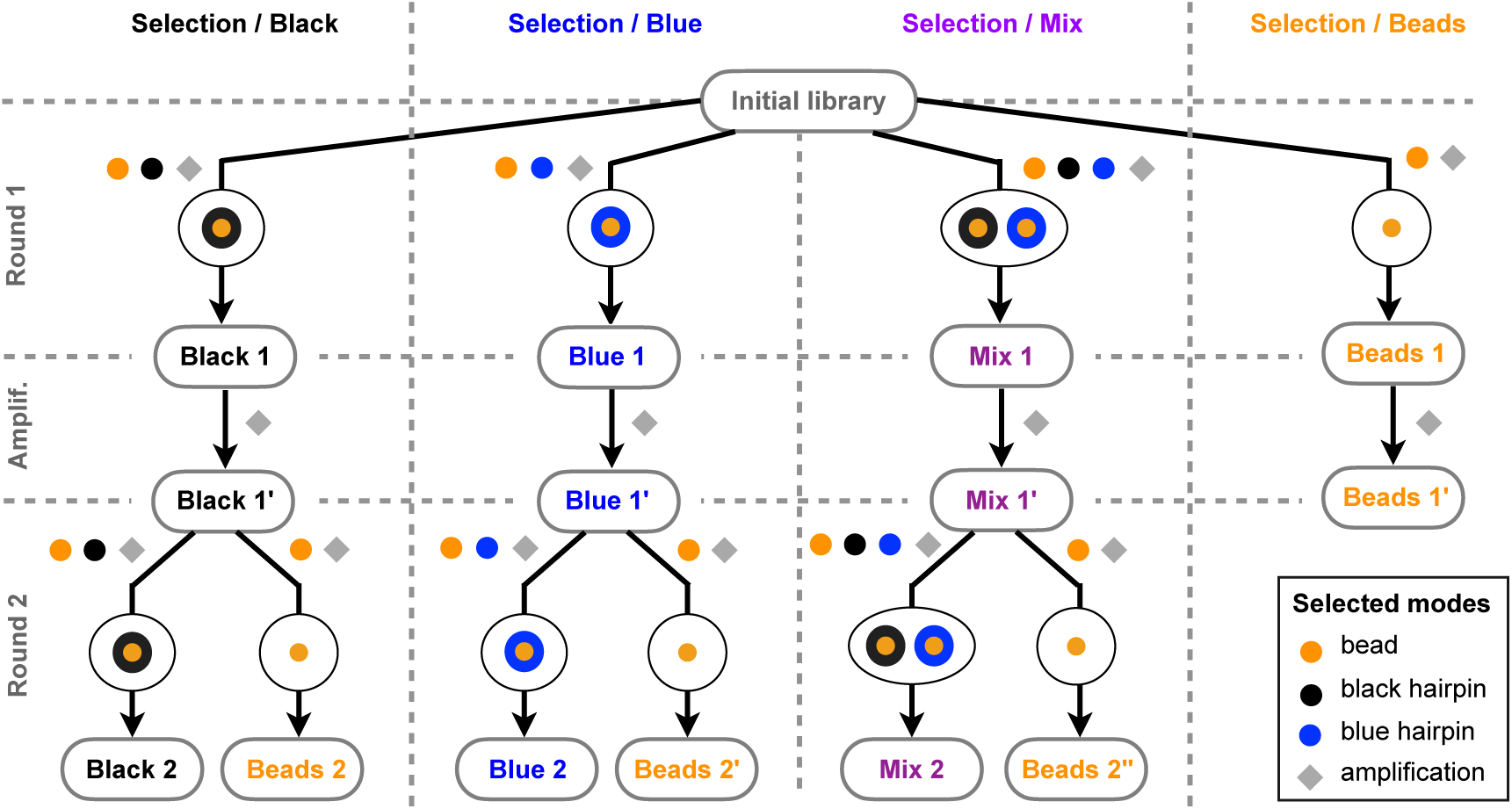
Training the model: selected set of modes. The selected modes incorporated in the model vary with each specific round. This adaptability enables comprehensive training using all available data. In the figure, the tree structure of the experiment (as presented in Figure 1 of the main text) is reported with the annotations of the selected mode for each branch or selection round. The four modes model distinct physical processes depicted in Fig. 5: binding the black hairpin, binding the blue hairpin, binding the supporting bead, and the bias introduced during the amplification process.

**FIG. 7.**
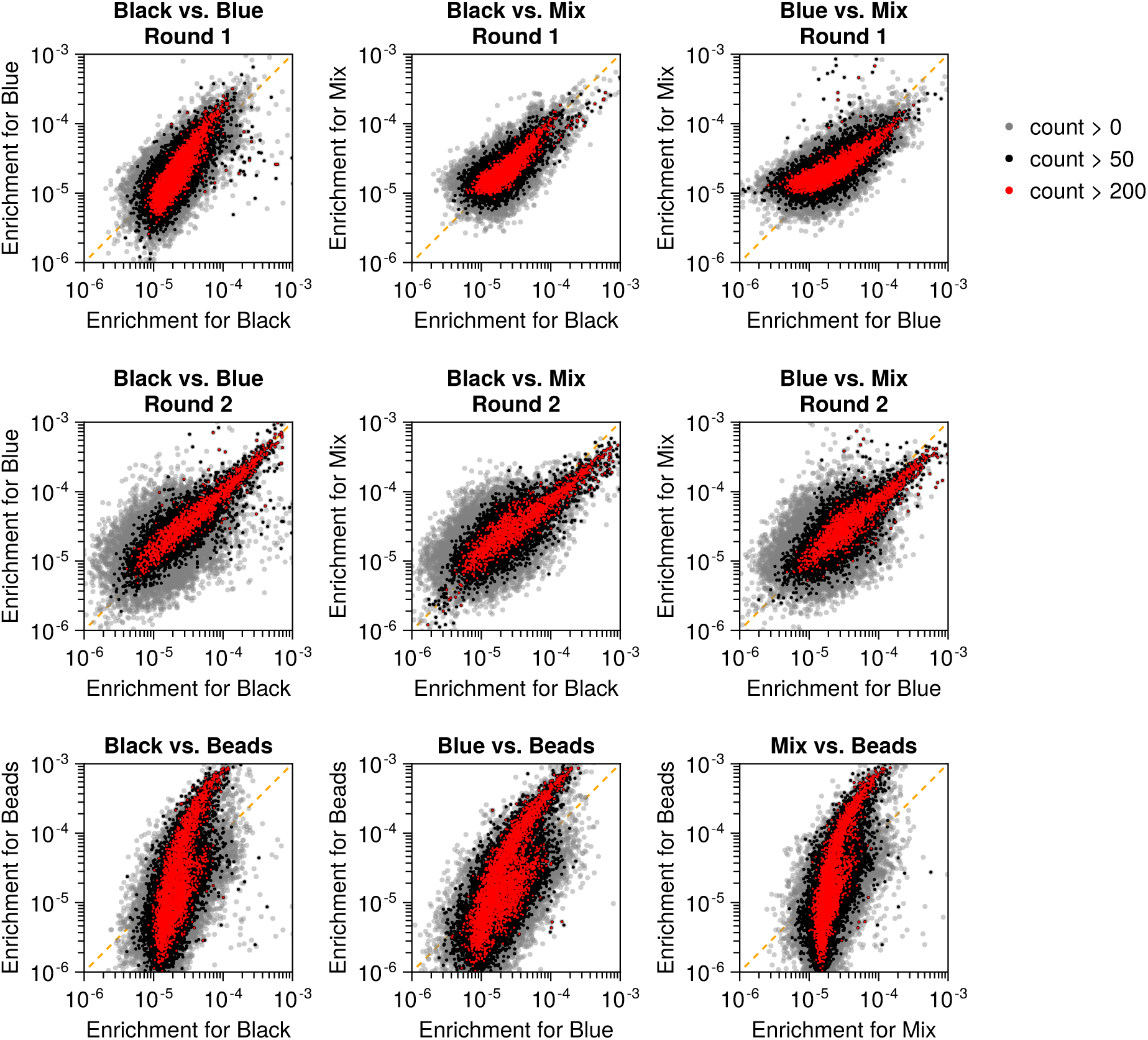
Comparison of empirical enrichments of each sequence in different experiments. The first and second row compare enrichments in the first and second rounds of selection (first and second row), against the Black, Blue, and Mix complexes. In each panel, the *x*-axis is the enrichment against a ligand (Black, Blue, or Mix), and the *y*-axis is the enrichment of the same sequence against a different ligand. The bottom row compares the enrichments against the Beads vs. the enrichments against the Black, Blue, and Mix ligands. See (B1) for the definition of enrichment. Sequences with counts lower than a given threshold are filtered out, as indicated in the legend at the top-right: all sequences with at least one read before and after selection are shown in gray, while sequences with more than 50 (200) reads before and after selection are shown in black (red). The Pearson correlations corresponding to each panel are shown in Fig. 8.

**FIG. 8.**
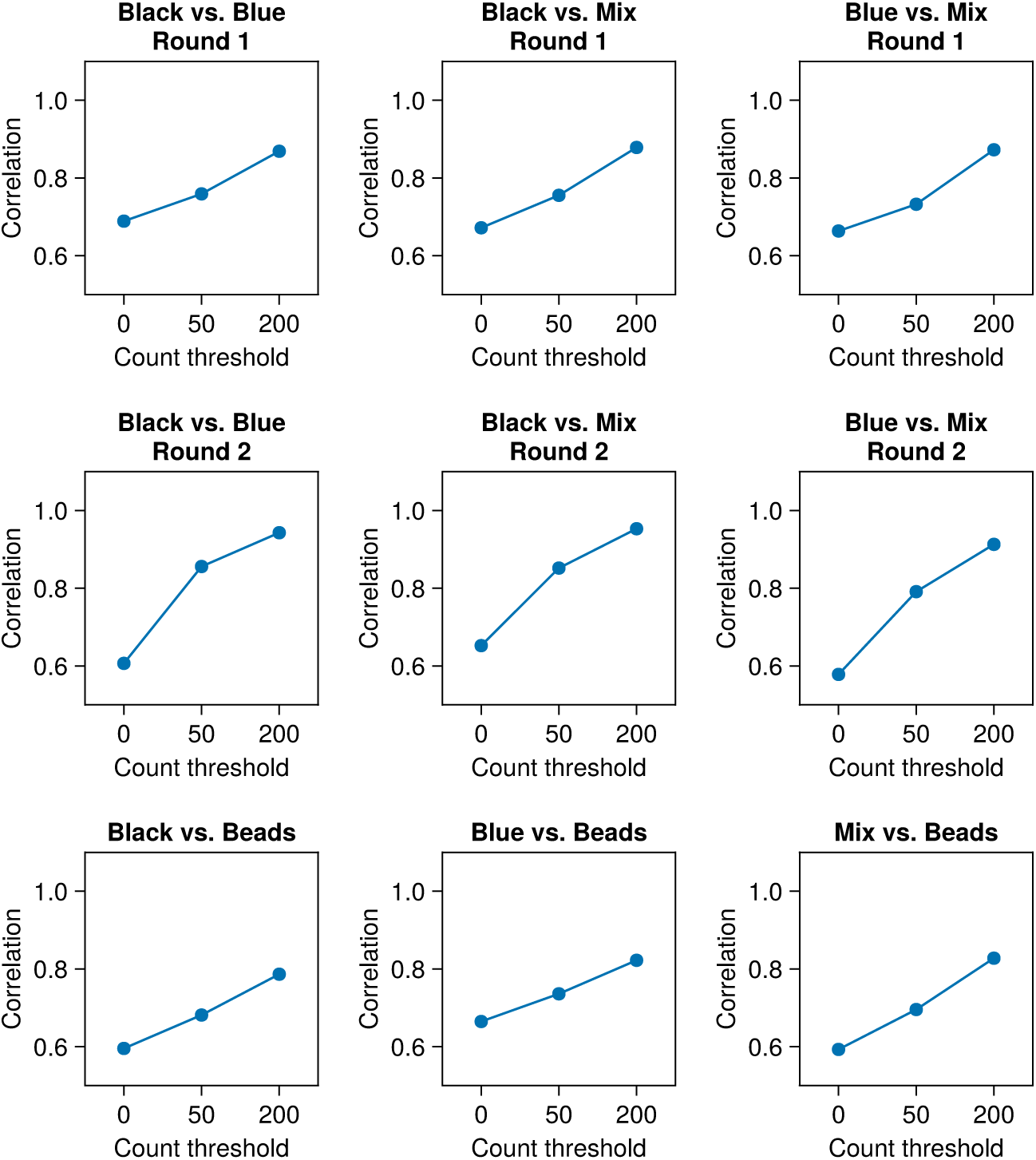
Pearson correlations between log-enrichments (log(*ϵ* _*st*_), where *ϵ* _*st*_ is defined in (B1)) in different experiments. Each panel shows the correlations between log(*ϵ* _*st*_) and log(*ϵ* _*st ′*_) for different experiments *t, t* ^*′*^, indicated in the panel title. The correlations are computed after filtering out sequences with counts lower than a given threshold (indicated in the *x*-axis) before and after selection. Panels are in correspondence to Fig. 7.

**FIG. 9.**
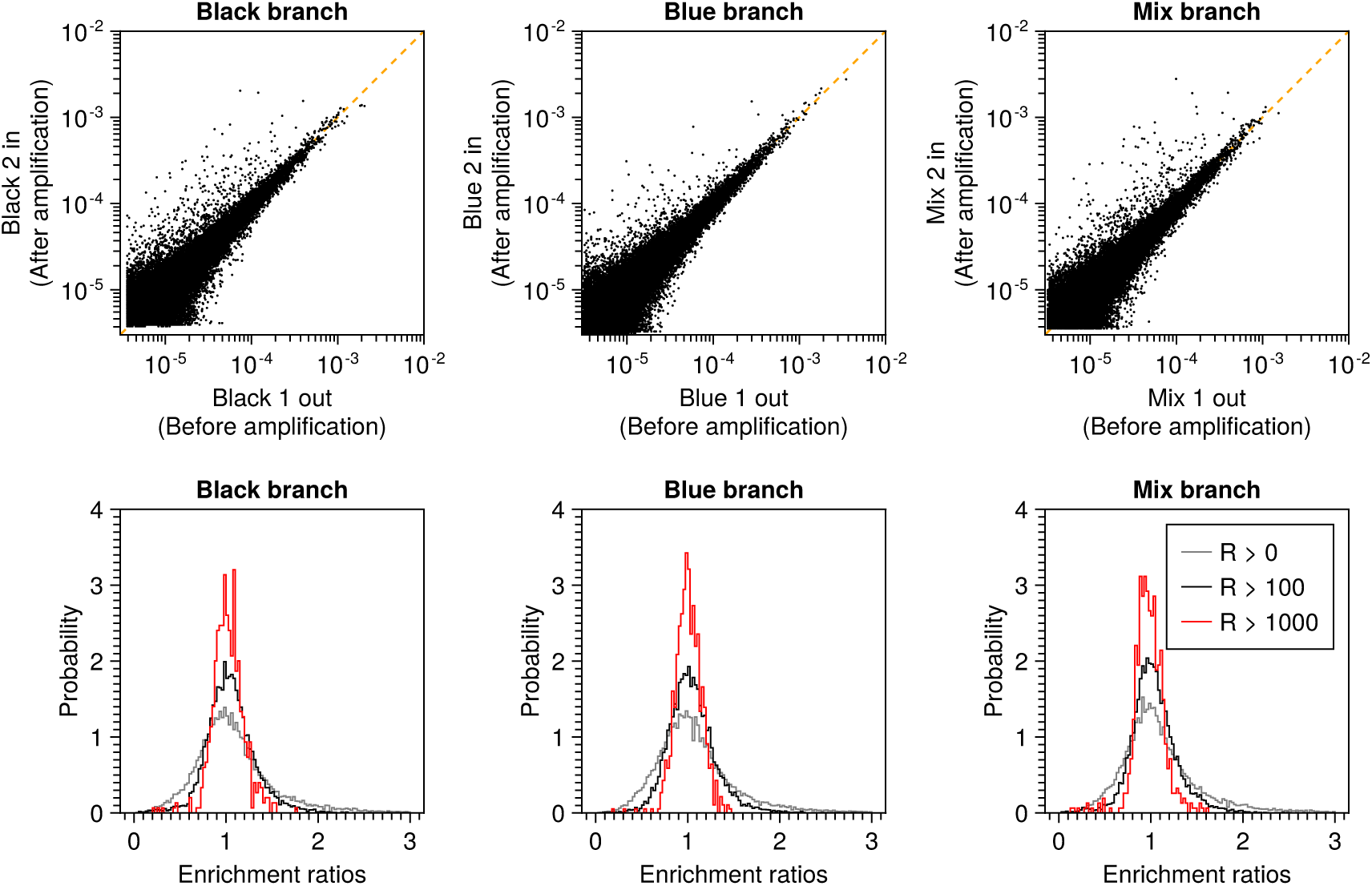
Sequencing reads are collected before and after amplification, after the first round of selection is finished and before starting the second round of selections. This was done for each branch of the experiment tree (see Fig. 1 in the main text): Black branch, Blue branch, and Mix branch. The top panels show a scatter of the normalized counts before (x-axis) and after (y-axis) amplification. The Pearson correlation coefficients are: 0.97 (for Black branch), 0.98 (for Blue branch), and 0.97 (for Mix branch). The bottom panels show histograms of the corresponding enrichment ratios (see (B1)), after filtering out sequences with counts below a threshold (indicated in the legend). The histograms concentrate around 1, indicating no selection.

**FIG. 10.**
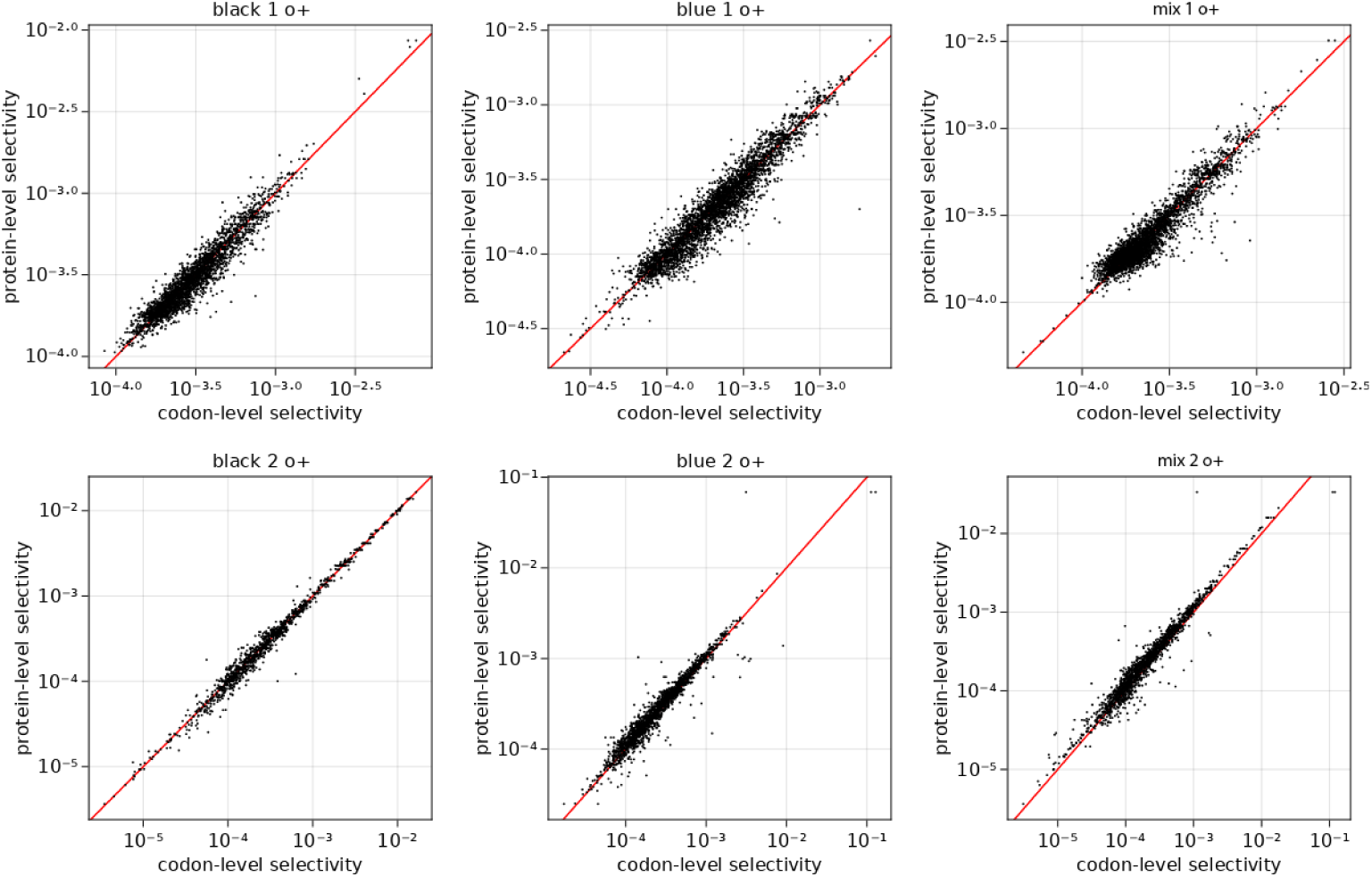
Each amino-acid sequence can correspond to many nucleotide variants due to codon degeneracy. The plots show a comparison of empirical enrichments ((B1)) for codon-sequences vs. the equivalent amino-acid sequences. Each panel corresponds to one experiment. Effects due to codon bias would be revealed by strong systematic dispersion in this plot. Sequences with less than 50 copies (at nucleotide or amino-acid level) are filtered out.

**FIG. 11.**
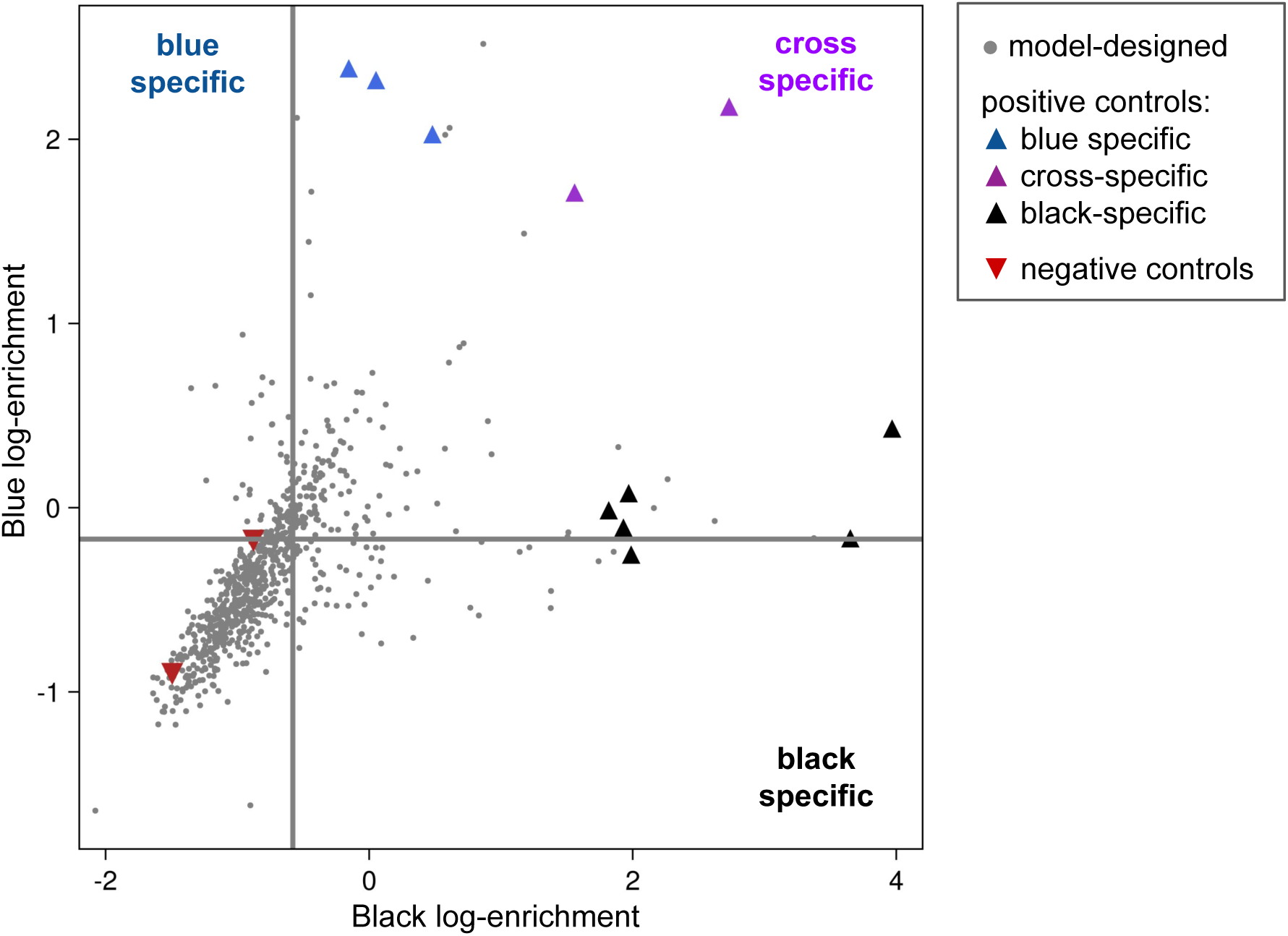
Experimental enrichment of model-designed sequences and controls. This figure illustrates the experimental enrichment results for model-generated sequences, marked in gray, and a selection of control sequences that serve as references and provide a sanity check for the experimental data. Among the selected sequences are the best specific binders, cross-specific binders, and two negatively selected sequences based on previous experiments. The horizontal lines represent the two predefined thresholds used to evaluate the generative performance, as detailed in Figure 3 of the main text. These thresholds correspond to the mean values of the two experimental enrichments. Notably, the negative and cross-specific control sequences align correctly with these thresholds. However, a majority of the specific sequences (blue and black triangles) fall outside the expected range. While more sophisticated methods for performance assessment exist, such as threshold selection based on blue-black enrichment ratios, we opted for a simpler approach in this instance. The primary goal here was to illustrate the model’s generative capacity, and a comprehensive definition of specificity falls outside the scope of this study.

**FIG. 12.**
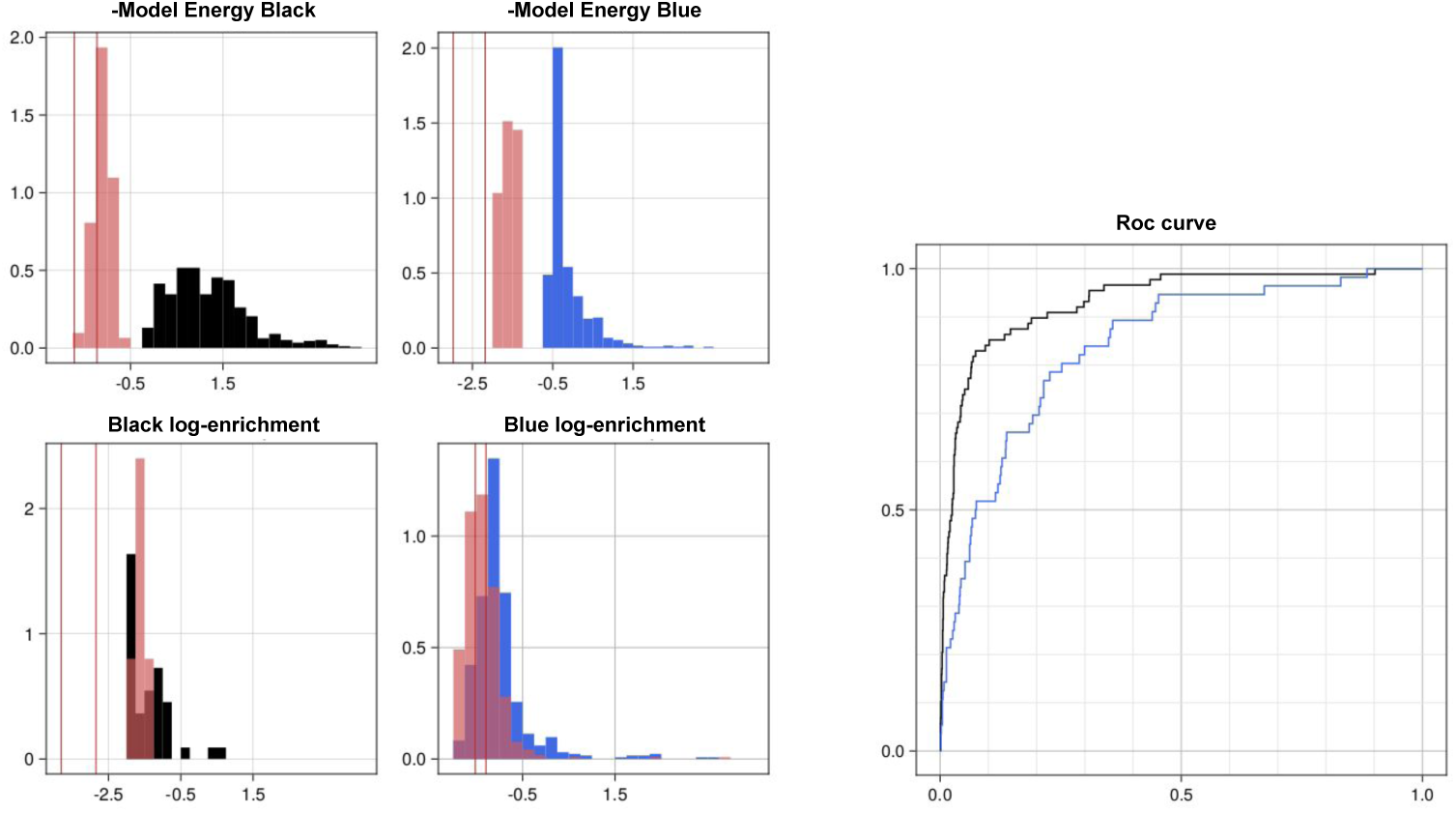
Validation of high-affinity antibodies generated via single-mode optimization. Designing high-affinity binders for a specific ligand, regardless of their affinity to other ligands, is a relatively easier task compared to selecting a specificity profile, as it involves no additional constraints on the optimization of single-model energies. In panels (a) and (b), the histograms depict the energies of designed sequences. Those in black and blue represent sequences predicted to exhibit high affinity for their respective ligands, while those in red are predicted to have low affinity. Red lines correspond to negative controls (see Fig. 11). Below these panels, two histograms illustrate the enrichment of these same sequences in validation experiments. Panel (e) showcases ROC curves for binder prediction using the model. Sequences above the threshold, denoted by the gray lines in panels (c) and (d), are considered binders. This threshold is set as the average of the enrichments plus one standard deviation. The model energy ranks these sequences, and the ROC curves are computed for both the black and blue ligand experiments. The Area Under the Curve (AUC) values are 0.92 for black and 0.79 for blue.

**FIG. 13.**
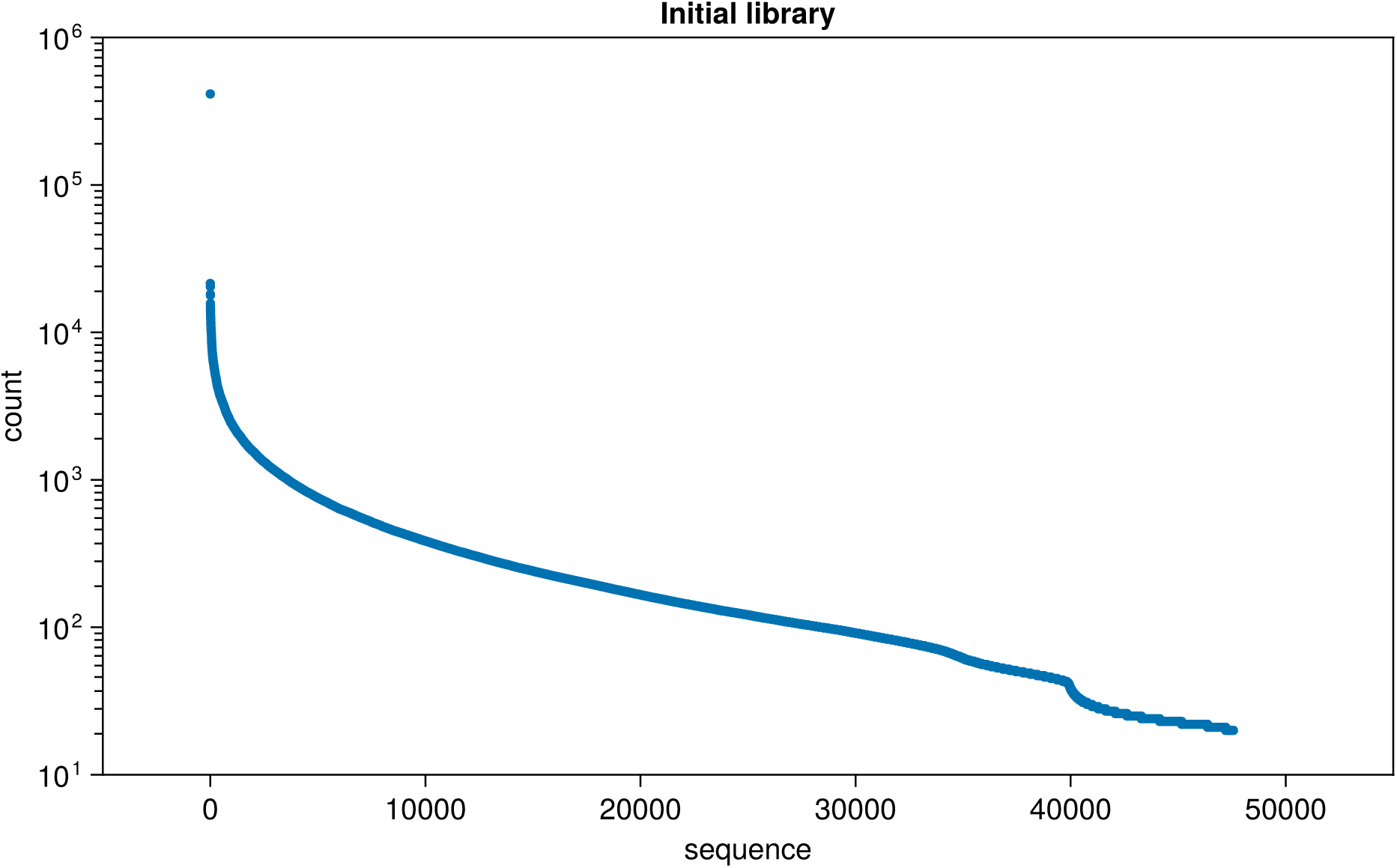
Number of reads of the initial library. Variants are displayed in descending order (from the most numerous variant to the less numerous one). The distribution is strongly non-uniform (note the log-scale on the y-axis).

**FIG. 14.**
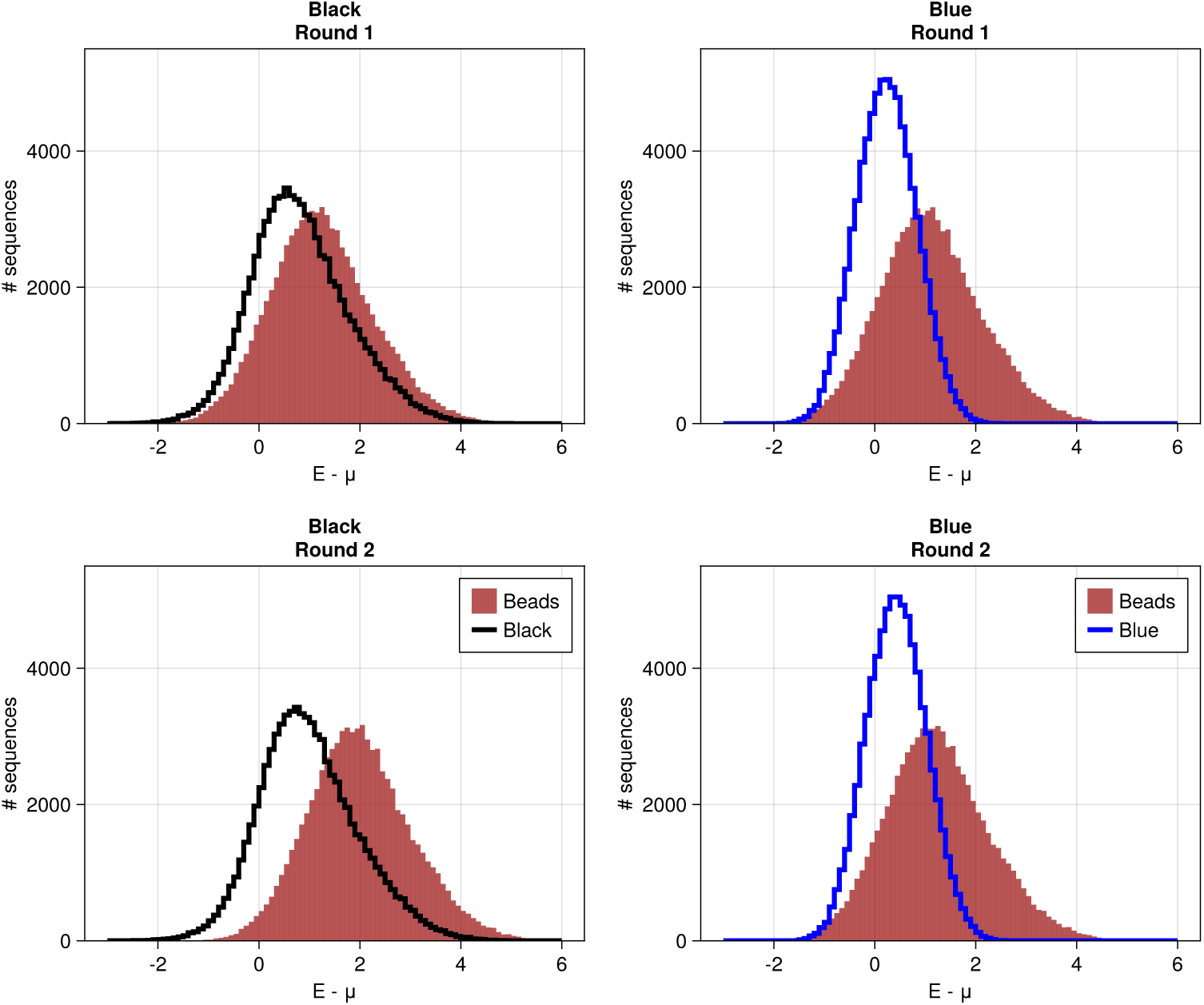
Histograms of *E*_*sw*_ − *μ*_*wt*_. The plots show the number of sequences *s* with a given value of *E*_*sw*_ − *μ*_*wt*_. The first column corresponds to experiments carried out with the Black target, and *w* is either the Black-bound mode (black line) or the Beads-bound mode (in brown). The right column corresponds to experiments carried out with the Blue target, and *w* is either the BLue-bound mode blue line) or the Beads-bound mode (in brown). The first and second rows correspond to the first and second rounds of selection. The binding energies to the Black and Blue targets are generally lower than the Beads binding energies.

## Notes

### Competing Interest Statement

The authors have declared no competing interest.

